# Level Dependent Subcortical EEG Responses to Continuous Speech

**DOI:** 10.1101/2024.04.01.587607

**Authors:** Joshua P. Kulasingham, Hamish Innes-Brown, Martin Enqvist, Emina Alickovic

**Affiliations:** Automatic Control, Department of Electrical Engineering, Linköping University, Sweden; Eriksholm Research Centre, Snekkersten, Denmark; Department of Health Technology, Technical University of Denmark, Lyngby, Denmark

## Abstract

The auditory brainstem response (ABR) is a measure of subcortical activity in response to auditory stimuli. The wave V peak of the ABR depends on stimulus intensity level, and has been widely used for clinical hearing assessment. Conventional methods to estimate the ABR average electroencephalography (EEG) responses to short unnatural stimuli such as clicks. Recent work has moved towards more ecologically relevant continuous speech stimuli using linear deconvolution models called Temporal Response Functions (TRFs). Investigating whether the TRF waveform changes with stimulus intensity is a crucial step towards the use of natural speech stimuli for hearing assessments involving subcortical responses. Here, we develop methods to estimate level-dependent subcortical TRFs using EEG data collected from 21 participants listening to continuous speech presented at 4 different intensity levels. We find that level-dependent changes can be detected in the wave V peak of the subcortical TRF for almost all participants, and are consistent with level-dependent changes in click-ABR wave V. We also investigate the most suitable peripheral auditory model to generate predictors for level-dependent subcortical TRFs and find that simple gammatone filterbanks perform the best. Additionally, around 6 minutes of data may be sufficient for detecting level-dependent effects and wave V peaks above the noise floor for speech segments with higher intensity. Finally, we show a proof-of-concept that level dependent subcortical TRFs can be detected even for the inherent intensity fluctuations in natural continuous speech.

**Visual abstract:** 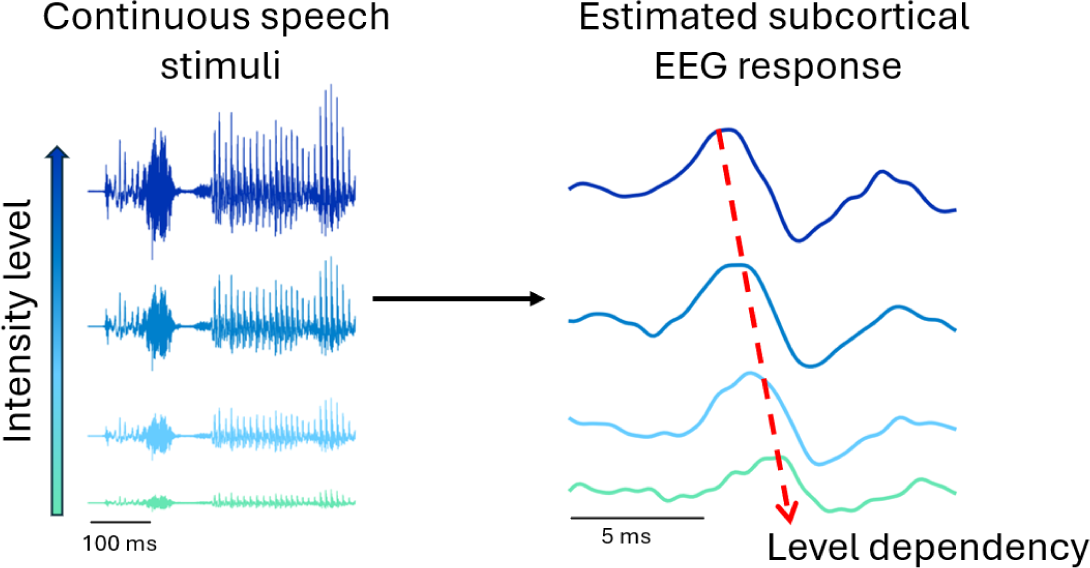

**Significance statement:** Subcortical EEG responses to sound depend on the stimulus intensity level and provide a window into the early human auditory pathway. However, current methods detect responses using unnatural transient stimuli such as clicks or chirps. We develop methods for detecting level-dependent responses to continuous speech stimuli, which is more ecologically relevant and may provide several advantages over transient stimuli. Critically, we find consistent patterns of level dependent subcortical responses to continuous speech at an individual level, that are directly comparable to those seen for conventional responses to click stimuli. Our work lays the foundation for the use of subcortical responses to natural speech stimuli in future applications such as clinical hearing assessment and hearing assistive technology.

## Introduction

The auditory brainstem response (ABR) is an electroencephalographic (EEG) measure of subcortical responses to auditory stimuli. It has been widely used to detect hearing impairments in newborns (Galambos & Despland, 1980; Picton et al., 1994) and coma patients (Guérit, 1999) for whom behavioral tests are not suitable. The ABR consists of distinct waveform peaks which correspond to one or more stages of the ascending auditory pathway. The latencies and amplitudes of these peaks depend on several characteristics of the stimulus such as frequency, intensity and presentation rate (Picton, 2010).

One shortcoming of conventional ABR measures is that several responses to short, repetitive stimuli need to be averaged in order to obtain ABR event related potentials (ERP). However, recent work has detected subcortical EEG responses to continuous, natural speech using linear deconvolution methods (Maddox & Lee, 2018), called the temporal response function (TRF). The TRF is a linear model that maps stimulus features (predictors) to the EEG in a continuous manner (Lalor et al., 2009). Although TRFs have been widely used to investigate *cortical* responses (Brodbeck & Simon, 2020; Kulasingham & Simon, 2023), there have been far fewer studies on *subcortical* TRFs (Polonenko & Maddox, 2021; Shan et al., 2024; Bachmann et al., 2021; Kulasingham et al., 2024).These subcortical TRFs have similar morphologies to conventional ABR ERPs, especially regarding the wave V peak (Bachmann et al., 2023). Moving towards ecologically relevant, natural speech stimuli could provide several benefits when investigating the auditory system (Hamilton & Huth, 2020), and may also increase patient comfort during clinical hearing assessments. Continuous speech is also better suited for studies involving hearing aid users, since hearing aids often employ speech-specific signal processing algorithms that are poorly suited for the transient stimuli used in conventional ERP methods (Laugesen et al., 2018).

A critical step towards using natural speech in such applications is determining whether subcortical TRFs display consistent level-dependent changes. It is reasonable to expect level-dependency in subcortical TRFs that is similar to level-dependency in ERPs to clicks or short speech syllables (Skoe & Kraus, 2010), since the underlying neural mechanisms are thought to be similar. However, there has been no experimental validation of this hypothesis, since prior studies on subcortical TRFs have not investigated changes in TRF morphology with intensity. There have, however, been a handful of studies that detected level-dependent changes in *cortical* TRFs(Drennan & Lalor, 2019; Verschueren et al., 2021; Van Hirtum et al., 2023).

Inspired by the need to validate the presence of level-dependency in subcortical TRFs, we investigate level-dependent changes in subcortical TRFs using EEG data collected from 21 participants listening to clicks and natural continuous speech stimuli with varying intensities. We estimate level-dependent TRFs by separating the stimulus into intensity bins and jointly estimating TRFs for each bin, following methods used for cortical level-dependent TRFs (Drennan & Lalor, 2019; Lindboom et al., 2023). This technique may reveal level-dependent patterns in the TRFs that can be compared with the level-dependency seen in conventional ABR ERPs.

A key consideration when deriving subcortical TRFs is the use of appropriate predictors. Since the TRF is a linear model, it cannot model the highly non-linear processing stages in the auditory periphery. Recent studies have used auditory peripheral models to generate predictors that incorporate these non-linearities (Shan et al., 2024; Kulasingham et al., 2024). These predictors are then used to fit TRFs, resulting in a more linearized estimation problem, and improved subcortical TRFs. However, some of these peripheral models also account for level-dependency effects, and it is unclear if predictors generated from these models would be suitable for detecting level-dependent changes in the subsequently estimated TRF waveform.

In this study, we explored the following research questions: 1) Are there level-dependent changes in subcortical TRFs. 2) Which predictor is most suitable for detecting level-dependent subcortical TRFs. 3) How consistent are level-dependent TRF patterns at an individual participant level. 4) How similar are level-dependent changes in TRFs compared to click ABR ERPs. 5) How much data is required to estimate level-dependent subcortical TRFs. 6) Can level dependent TRF changes be detected using both short (5 second) and long (1 minute) periods of fixed stimulus intensity. 7) Can level-dependent TRFs be detected for inherent variations of stimulus intensity in natural speech. This work lays the foundation for the use level-dependent changes in subcortical TRFs for applications such as detecting hearing thresholds, evaluating hearing aid benefits and investigating hearing impairments using natural continuous speech stimuli.

## Materials and methods

### Experimental details

EEG data were collected from 24 native Danish speaking participants (mean age = 32.4, standard deviation (SD) = 8.8 years, 12 male, 12 female). The study was approved by the ethics committee for the capital region of Denmark (journal number 22010204) and the Swedish Ethical Review Authority (number 2023-03074-01), and all participants provided written informed consent. A pre-screening audiogram was conducted, and 22 participants had hearing thresholds below 30 dBHL from 250 Hz - 4 kHz in both ears. The other 2 participants were excluded from the analysis due to having hearing thresholds above 30 dBHL at 8 kHz. Another participant did not have clear wave V peaks in the click ERP or the TRF and was excluded from further analyses, but is shown in the supplementary figures for completeness. Therefore, the data from 21 participants were used for all further analyses. The participants were seated listening to click trains or continuous speech excerpts from a Danish audiobook (Simon, a biography on Simon Spies, read by a male speaker) while their EEG data was collected.

The stimuli were presented binaurally using an RME Fireface UCX soundcard (RME Audio, Haimhausen Germany) and Etymotic ER-2 (Etymotic Research, Illinois, USA) insert earphones, which were shielded using a grounded metal box to avoid direct stimulus artifacts on the EEG. All stimuli were presented at 44.1 kHz. The participants were asked to listen to the clicks while keeping their eyes fixated on a cross presented on a screen that was placed 1.5 meters in front of them.

The experiment consisted of two conditions, a click stimulus condition and a speech stimulus condition. The stimuli for both conditions were presented at 4 different intensity levels. The click stimuli consisted of alternating rarefaction and condensation clicks at a rate of 44 clicks per second. The clicks were rectangular with 91 µs duration (4 samples), and were distributed according to a pseudo-Poisson distribution with a minimum inter-click-interval of 15 ms. The loudest stimulus was calibrated to be 66 dBA using a measurement amplifier (Bruel and Kjær Type 2636, with a time averaged measurement of 1 second), an inner ear simulator (Bruel and Kjær Type 4157) and a Bruel and Kjær Type 4231 sound source. Although this method is not typically used for click intensity calibration, we chose to use this because the same method was used for the speech stimuli, allowing for a consistent calibration pipeline, and because the absolute intensity level of the clicks are not as important to this study as the relative differences between intensity levels, which were constructed by scaling the stimulus files. The clicks at 66 dBA were perceptually quite loud, but not uncomfortable, based on self reports of the participants. 3 other click levels were also constructed by downscaling the click stimulus amplitudes to result in 60, 48 and 36 dBA. These 4 levels were presented at the start of the experiment, with 2 minutes in each intensity (5280 clicks), in decreasing order of intensity, with short 5 second breaks in between.

After the click condition, the speech condition was presented, which had 80 trials, each consisting of 1 minute excerpts from the audiobook, where prolonged silent periods were shortened to a maximum of 500 ms. The speech was calibrated using the same measurement amplifier as the clicks, with the loudest intensity level being 72 dBA. 4 other intensities were also constructed by downscaling the stimulus amplitudes to result in 60, 48 and 36 dBA stimuli. The speech consisted of two types of trials: fixed intensity for a long duration (1 minute) or a short duration (5 seconds). For the long duration type, the intensity for the entire 1-minute trial was fixed to the same intensity value (72, 60, 48 or 36 dBA). For the short duration type, the intensity within the 1-minute trial changed randomly across the 4 intensity levels every 5 seconds, with a cosine ramp of width 500 ms in between the level changes, such that each level was presented the same amount of times (i.e., 3 times for each of the 4 levels in a 60-second trial). These two types of trials were employed to investigate whether level dependent responses could be detected to both long and short periods of fixed intensities. 40 trials of each type were fully intermixed randomly, for a total of 80 trials (80 minutes), and presented to the participants. The narrative order of the audiobook story was preserved, and only the intensity levels and trial types were randomized. There was a short (∼ 5 second) break after each trial. The participants were asked to follow the story while looking at a fixation cross and respond to a comprehension question that was presented on a screen placed 1.5 meters in front of them. The question was presented after a random number of trials, distributed uniformly between 3 and 6 trials. Participants were required to respond by pressing a button on a keyboard in front of them, with options to choose either YES, NO or DON’T KNOW. These questions were presented only to encourage participants to focus on the story, as commonly done in cortical TRF studies, and were not further analyzed. After each block of 20 trials, the participants could take a longer break from the experiment. In summary, the entire speech condition consisted of 80 minutes of speech stimuli (40 long duration, 40 short duration fixed intensity) presented at 4 intensity levels.

### EEG preprocessing

A Biosemi Active 2 system (Biosemi, Amsterdam, The Netherlands) was used to collect EEG data using a 32 channel cap with electrodes in standard 10-20 positions. Electrodes were also placed on both mastoids, earlobes and above and below the right eye. The data were collected at a sampling frequency of 8192 Hz with a fifth order cascaded integrator-comb (CIC) antialiasing filter with a -3 dB point at 1638.4 Hz. The data was further filtered offline with a delay compensated FIR filter with a cutoff frequency of 1 Hz. All data analysis was conducted using MATLAB (version R2021a) and the Eelbrain Python toolbox (version 0.38.1) (Brodbeck et al., 2021) on a laptop with an AMD Ryzen 7 CPU with 32 GB RAM. Only the Cz channel, referenced to the average of the two mastoid channels, was used for this subcortical study to be analogous to commonly used EEG montages for click-ABR measurements. Power line noise was removed using FIR notch filters with widths of 5 Hz and center frequencies at all multiples of 50 Hz up to 1000 Hz. Simple artifact rejection was performed by detecting segments of the EEG data that were 5 standard deviations (SD) above or below the mean and setting 1 second sections around these points to zero in both the EEG and the predictors used for TRF analysis (∼ 0.5 − 4.8% data excluded per participant). Since detecting subcortical responses requires millisecond resolution, the audio output of the soundcard was also fed to the Erg1 channel of the biosemi EEG system via an optical isolator (StimTrak, BrainProducts, GmBH, Gilching, Germany), to avoid timing jitters or clock drifts between the stimulus presentation system and the EEG measurement system (Kulasingham & Simon, 2023). These Erg1 signals were later used to align the stimulus waveforms for TRF analysis.

### Click ERP estimation

Click evoked ABRs were estimated for each of the four intensity levels separately. The EEG data was filtered between 30 and 1000 Hz using a delay compensated FIR bandpass filter. Click onsets were detected using the recorded stimulus on the Erg1 channel and EEG epochs from -10 to 30 ms around each click were extracted (5280 epochs in total). Epochs for both rarefaction and condensation clicks were then averaged together, following the standard procedure for click ERPs. Finally, the ERPs were baseline corrected by subtracting the average voltage value from -10 to 0 ms.

### Speech predictors

The speech stimuli were used to compute several types of predictors. These predictors approximate the non-linear processing stages of the peripheral auditory system, at differing levels of complexity. A detailed description of the predictors are presented in a recent work (Kulasingham et al., 2024), and is briefly recounted here. Five types of predictors were generated as given below, and the lags inherent in the output of each model were accounted for by shifting the generated predictors to align with the rectified speech predictor.

### Rectified speech predictor (RS)

The RS predictor was formed by half-wave rectifying the speech stimulus, following prior work on subcortical TRFs (Maddox & Lee, 2018). This serves as a coarse approximation of the rectifying properties of the peripheral auditory system.

### Gammatone spectrogram predictor (GT)

The speech was passed through a gammatone filterbank consisting of 31 filters from 80-8000 Hz with 1 equivalent rectangular bandwith (ERB) spacing as implemented in the Auditory Modelling Toolbox (AMT, version 1.1.0 (Majdak et al., 2022), function auditoryfilterbank with default parameters). The resulting amplitude spectra were averaged over all bands to form the predictor.

### Osses model predictor (OSS)

The speech stimuli were passed to the auditory model provided in (Osses Vecchi & Kohlrausch, 2021), which is based on the model in (Dau et al., 1996). The implementation in AMT was used (function osses2021 with default parameters). Only the initial stages of this model were used (outer ear pre-filter and gammatone filterbank followed by rectification and lowpass filtering), as fully described in a prior work (Kulasingham et al., 2024). The resulting signals from 31 center frequencies were averaged together to form the predictor.

### Osses model predictor with adaptation (OSSA)

The next stage of the Osses model, which includes adaptation loops (Osses Vecchi & Kohlrausch, 2021) were now included in addition to the above-mentioned stages. The 31 channel output from the adaptation loops were averaged together to generate the predictor. Notably, unlike the previous models, this model accounts for the level-dependent adaptation properties of the early auditory system to some extent.

### Zilany model predictor (ZIL)

Finally, a more complex auditory model (Zilany et al., 2014) was used to generate predictors. This model consists of several stages approximating non-linear cochlear filters, inner and outer hair cell properties, auditory nerve synapses, and adaptation. The implementation in the Python cochlea package (Rudnicki et al., 2015) was used with 43 auditory nerve fibers with high spontaneous firing rates and center frequencies logarithmically spaced between 125 Hz and 16 kHz, in line with previous work (Shan et al., 2024). The outputs of this model are the mean firing rates of the auditory nerves, which were averaged to form the final predictors. It is noteworthy that this model also accounts for the level-dependent properties of the early auditory system, probably more accurately than the OSSA model, due to its more complex and biophysically inspired structure.

### Level Dependent Temporal Response Function Estimation

The simple TRF model can be described using a regression model

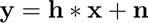

where **y** ∈ ℝ*^N^* denotes the vector of EEG measurements, **h** ∈ ℝ*^K^*denotes the TRF vector (impulse response), **x** ∈ ℝ*^N^* denotes the predictor, ∗ denotes the convolution operation, and **n** ∈ ℝ*^N^*denotes the noise.

To investigate level-dependent TRFs, the predictor was separated into equal intensity bins, based on the presented intensity (72, 60, 48 or 36 dBA), following the method outlined for level-dependent cortical TRFs in previous cortical studies (Drennan & Lalor, 2019; Lindboom et al., 2023). For each intensity level, a new predictor was constructed by setting to zero the predictor time segments that were not presented at that intensity. This resulted in 4 predictors, one for each intensity bin. These 4 predictors were then normalized to allow for all intensity levels to contribute equally to the TRF model. The EEG data was filtered between 30 and 1000 Hz using a delay compensated FIR bandpass filter. The EEG and the 4 predictors were then used in a joint regression model

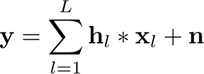

to estimate TRFs. Here the sum is over the number of intensity levels (*L* = 4) with **h***_l_* ∈ ℝ*^K^* denoting the TRF vector (impulse response) for level *l*, and **x***_l_* ∈ ℝ*^N^* denoting the constructed predictor for level *l*. This regression can be solved using the least squares method for linear regression, by constructing a design matrix with shifted versions of the predictor (Alickovic et al., 2019; Crosse et al., 2021). The solution is given by

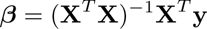

where ***β*** ∈ ℝ*^KL^* is the concatenated TRFs for all levels and **X** ∈ ℝ*^N×KL^* is the concatenated design matrix of shifted versions of all predictors. This method is very similar to the commonly used Ridge regression approach for cortical TRFs (Crosse et al., 2021). However, for this work, we do not use any regularization, since we found that unregularized regression provided clear TRFs without the added complexities of optimizing the regularization parameters or disentangling confounds due to regularization in level-dependent TRFs. Using unregularized TRFs is also consistent with several earlier studies on subcortical TRFs (Maddox & Lee, 2018; Polonenko & Maddox, 2021; Shan et al., 2024; Kulasingham et al., 2024; Bachmann et al., 2023).

This method results in a TRF for each intensity level, allowing for the investigation of level-dependent effects on TRF waveforms. Upon initial visual inspection, the wave V peak was the only clearly visible feature in the TRFs (similar to prior work (Maddox & Lee, 2018; Bachmann et al., 2021; Kulasingham & Simon, 2023)), and hence a smoothing was applied to the TRFs using a Hamming window of width 4 ms that was moved over every sample. This helped to eliminate high frequency noise and led to cleaner wave V estimates, especially since some of the TRFs were more noisy due to each intensity level having only 1/4^th^ of the data. This same smoothing was also applied to the click ERPs to have a consistent pipeline and facilitate fair comparison. The TRFs were then scaled to have the same root mean square (r.m.s.) amplitude as the average click ERPs, using a common scaling factor for all participants, to enable morphology comparison with the click ERPs.

Note that although the entire dataset has 80 minutes of data in total, the TRF for each level was fit using binned predictors that had only 20 minutes of non-zero data. This dataset was then further divided into the long and short fixed intensity duration conditions (40 minutes in each condition with 10 minutes of data at each level), which resulted in noisier estimates. To investigate the effect of data length on TRF features, portions of the 40 minutes of data from the short duration condition were used. The portions consisted of data in increments of 4 minutes up to 40 minutes (or 1-minute increments per level).

### Level dependent TRFs for inherent intensity fluctuations

Finally, a proof-of-concept for detecting level dependent TRFs to the inherent changes of intensity in natural speech was investigated. The intensity bins were determined using the smoothed predictor waveform instead of the ground-truth intensities of the presented stimuli. The predictor was smoothed using a 300 ms Hamming window to approximate a dynamic measure of inherent intensity (see Discussion for comments on alternative measures of intensity). For this analysis, the number of intensity bins was increased to 8 to investigate whether a finer resolution of level-dependent changes could be detected. The intensity bin edges were determined as 8 values that segmented this signal into an equal amount of data within each bin. That is, the 1/8^th^ percentiles of this smoothed predictor were used as the values by which to segment the predictor into intensity bins. The original predictor was separated into intensity levels using these segments, and 8 new binned predictors for each level were constructed as detailed above. These 8 predictors were used to jointly fit TRFs following the same least squares procedure as before.

We preferred to construct the intensity bins to have an equal amount of data in each bin, rather than to have intensity bins that were equally spaced logarithmically. This would eliminate possible confounds due to varying SNR when fitting TRFs with different amounts of data in each bin. It is noteworthy that in the previous analyses using the ground-truth intensity bins, the 4 bins were both equally spaced in intensity *and* had equal amounts of data by experimental construction. Additionally, since the smoothed predictor itself was used to determine the bins, there is the possibility that a low intensity section of a trial presented at a high intensity (e.g., a soft section of the 72 dBA trial) would be assigned to a lower intensity bin (and vice versa). In the previous version with the ground-truth intensities, the entire trial presented at a high intensity would be assigned to the high intensity bin (and vice versa). Therefore, this method using the smoothed predictor to split the data into intensity bins may provide a more accurate separation based on the inherent intensity levels that were actually heard.

### Performance metrics

The wave V peak of the subcortical TRFs was extracted by detecting the largest peak between 4-10 ms. The SNR of the peak amplitude was calculated as SNR = 10 log_10_((*S* − *N*)*/N*) where the power in a 5 ms window around the wave V peak was used as the measure of signal power *S* and the noise power *N* was estimated as the average power from -10 to 0 ms of the TRF. An SNR of 0 dB corresponded with visually distinct peaks and was set as the threshold for a meaningful wave V peak (signal power is equal to the noise power), similar to prior work (Polonenko & Maddox, 2021; Bachmann et al., 2023). For visualization purposes, the SNR was set to have an arbitrary minimum of -5 dB, since any SNR below 0 dB was considered to indicate the lack of a clear peak.

### Statistical analysis

The wave V peak amplitudes and latencies of the click ERPs and the speech GT TRFs were correlated across participants. To further compare the individual level-dependent patterns across click ERPs and speech GT TRFs, a simple linear fit was estimated for each participant’s amplitude and latency across the 4 intensity levels. The slopes and intercepts of these linear fits for click ERPs and speech GT TRFs were then correlated across participants. Pearson’s correlations were used with p-values corrected for multiple comparisons using the Holm-Sidak method. Similar linear fits were also performed for different datalengths and the slopes were investigated as a measure of the level-dependency of the TRFs.

## Results

### Level dependent subcortical responses for click and speech stimuli

EEG data from 21 participants listening to clicks and continuous speech at different intensities were analyzed using the midline central Cz channel, referenced to the average of the two mastoid electrodes. ERPs and subcortical TRFs were estimated for the click and speech stimuli, respectively. The average click ERPs and speech TRFs across participants are shown in Fig. 1. The click stimuli were presented at 66, 60, 48 and 36 dBA (see Methods). The click ERPs were estimated separately for each level and show clear wave V peaks that change with the intensity level of the stimulus (see also supplementary Fig. S1 for individual click ERPs). As expected, the wave V peak amplitude increases and the latency decreases with increasing intensity. Earlier peaks of the canonical ABR are not visible, similar to some prior studies (Maddox & Lee, 2018; Shan et al., 2024; Bachmann et al., 2023).

**Figure 1:**
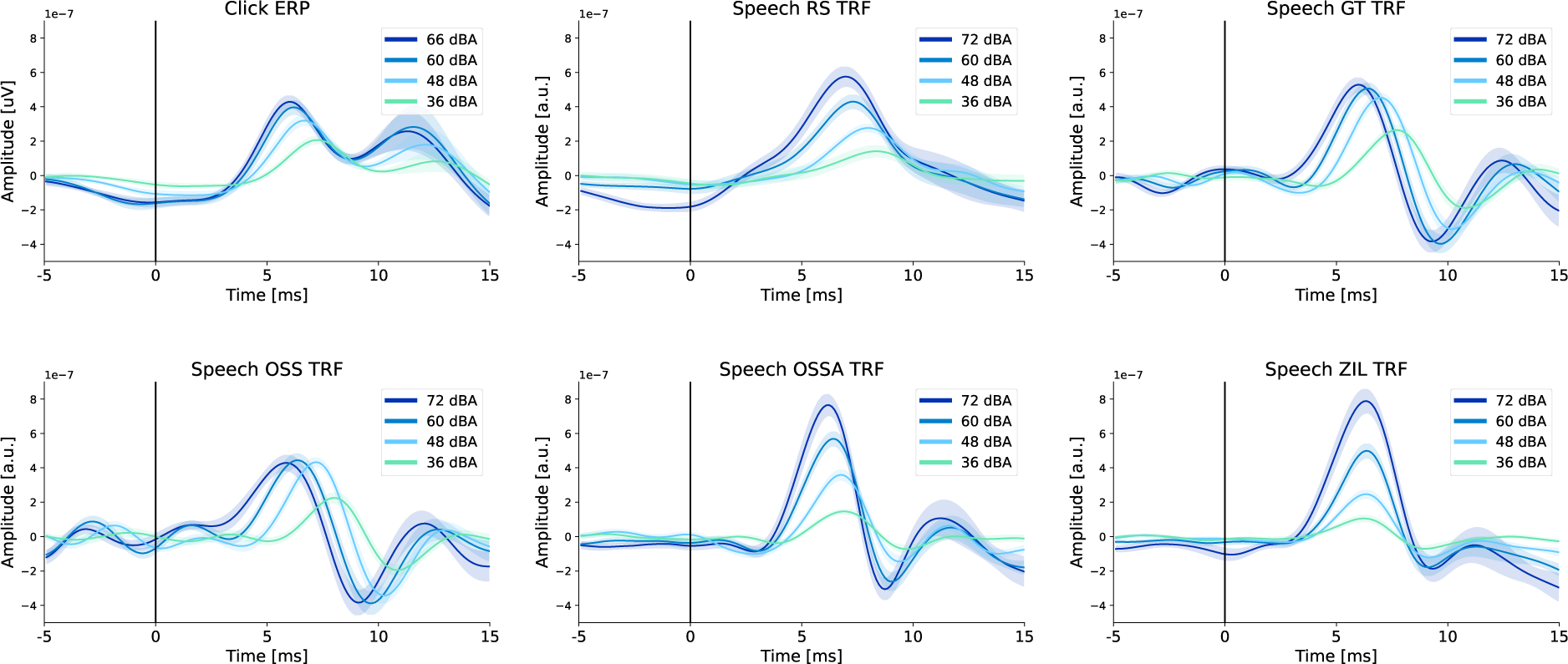
Average level-dependent click ERPs and speech TRFs. The average click ERP and speech TRFs (for the 5 predictors) across 21 participants are shown for each intensity level, with shaded areas representing standard error of the mean (s.e.m.). Clear level-dependent changes in wave V peaks are seen in all cases. Note that the ZIL TRF does not show a wave V latency change, since the ZIL model accounts for the non-linear latency effects in the predictor before it was used to fit TRFs. The OSSA TRF shows a reduced latency effect, since the OSSA model also accounts for level-dependent effects to some extent.

The speech TRFs were estimated using the 5 types of predictors (RS, GT, OSS, OSSA and ZIL) from EEG data recorded while listening to continuous speech from an audiobook. The speech was presented at a constant intensity of 72, 60, 48, or 36 dBA for either a short (5 seconds) or long (1 minute) period before changing randomly to another of the 4 intensity levels (see Methods for a full description of the stimuli). For this initial analysis, 40 minutes of the short condition and 40 minutes of the long condition were combined, and TRFs were estimated using unregularized regression (see Methods for more details). While the simple RS predictor showed level-dependent wave V peaks, it was notably noisy in many participants (see Fig. 2, and supplementary Fig. S2 with all individual TRFs), as expected from prior work (Kulasingham et al., 2024; Bachmann et al., 2023; Shan et al., 2024). The average TRFs for both GT and OSS predictors showed clear level dependency in both the wave V peak amplitude and the latency.

**Figure 2:**
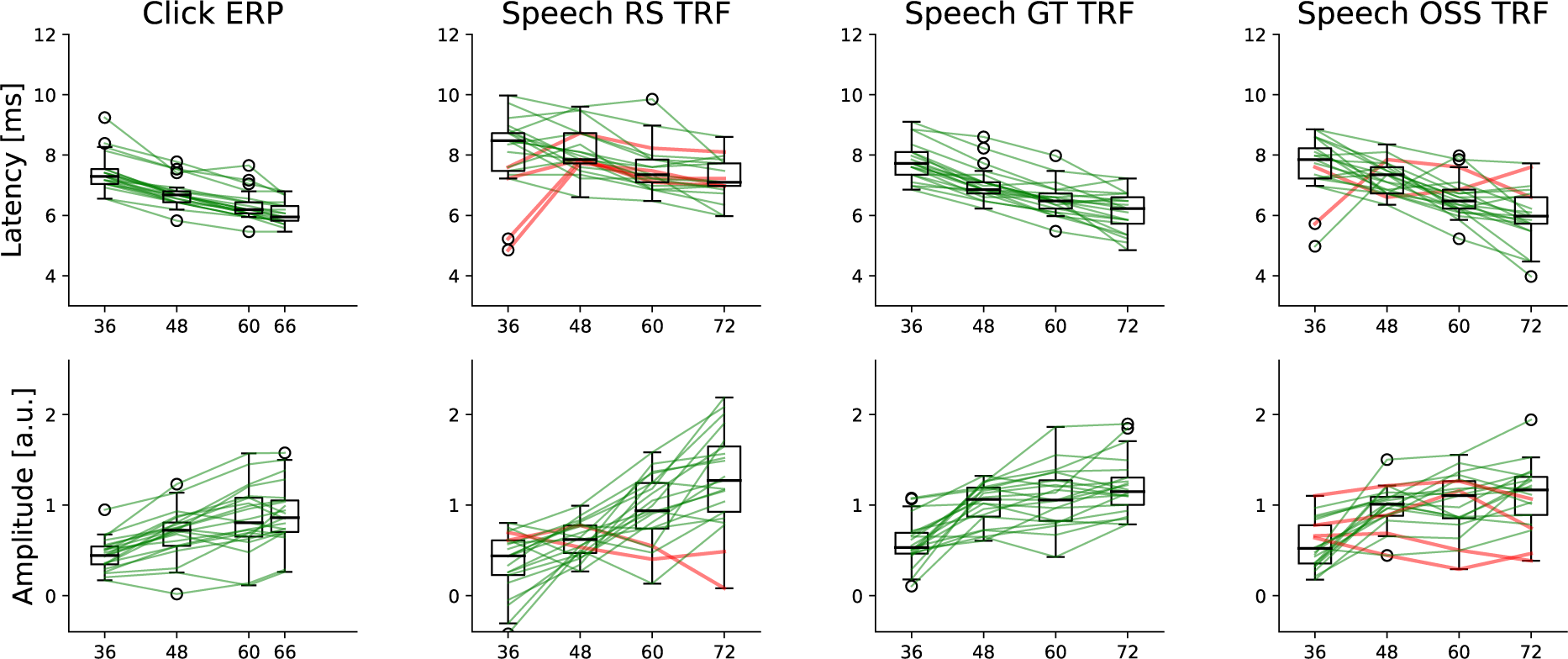
Individual changes in wave V peak amplitudes and latencies. The wave V changes for the click ERPs and subcortical TRFs are shown in each column. The OSSA and ZIL TRFs are omitted since they do not show clear latency changes. **Top row**: Latency changes, with boxplots across participants and each line denoting individual participants. Participants that have a smaller latency for the highest intensity compared to the lowest intensity (i.e., the expected trend) are shown in green and others are shown in red. **Bottom row**: Amplitude changes for each intensity. Participants that have a larger amplitude for the highest intensity compared to the lowest intensity (i.e., the expected trend) are shown in green and the others are shown in red.

Interestingly, the OSSA TRF showed a reduced peak latency change with level, and the ZIL TRF showed no level-dependent peak latency changes. However, both OSSA and ZIL showed level-dependent peak amplitude changes. The lack of clear peak latency changes can be explained by the fact that both OSSA and ZIL models account for the non-linear level-dependent processing of the auditory system using adaptation models. This results in the OSSA and ZIL outputs being ‘corrected’ for level-dependent effects. Therefore, using OSSA and ZIL outputs as predictor inputs to the TRF model results in a lack of level-dependent latency effects in the TRF waveform. The slight latency change seen in the OSSA model implies that the OSSA model is not as accurate in accounting for level-dependent effects as the ZIL model. The peak amplitude change in the OSSA and ZIL TRFs is due to the normalization of the predictor within each level before fitting TRFs. This normalization results in an upscaling of the lower intensity predictors, which leads to a corresponding downscaling of the fitted TRFs. This predictor normalization is useful since it allows for all intensity levels to drive the TRF estimation equally, as reported in prior work on level-dependent cortical TRFs (Drennan & Lalor, 2019). In addition, normalization would allow for clear detection of level-dependent effects in the TRFs for all predictors, since the input predictors at each intensity level would be at the same scale.

Accounting for level-dependency within the predictor results in a more ‘linearized’ input-output system for the TRF, leading to TRF models with larger wave V peaks (see (Shan et al., 2024; Kulasingham & Simon, 2023)). However, for our purposes, this is unsuitable since accounting for level-dependent effects in the predictor leads to a lack of clear latency changes in the TRF. Therefore, even though the OSSA and ZIL models are more complex models of the auditory periphery that incorporate more non-linearities, we reach the conclusion that the simpler GT and OSS models are more suitable for the purpose of this study, which is to detect clear level-dependent changes in the TRF waveform that are analogous to the level-dependent changes seen in the click ERPs. Given that these level-dependent changes most reliably manifest in the latency of the responses, it is important not to account for latency differences in the predictor used to estimate the responses.

### Level-dependent changes in wave V peaks are consistent at the individual level

The wave V peak amplitudes and latencies were extracted for both click ERPs and speech TRFs as the peak between 4-10 ms. The wave V amplitudes and latencies are shown for each level across participants in Fig. 2. The OSSA and ZIL TRFs are not shown since they do not show clear latency effects and were not investigated further. For the click ERPs, the amplitudes increase and latencies decrease with increasing intensity in a monotonic fashion for almost all participants. However, the individual trends for the speech TRFs were considerably noisier. To simplify comparisons between the predictors, the lowest (36 dBA) and highest (72 dBA) intensity levels were considered (also shown in Fig. 2). The RS and OSS TRFs appear noisier, with some participants not showing the expected level-dependent amplitude and latency change. The number of participants with level-dependent changes that were against the expected trend (red lines in Fig. 2) were as follows: RS latency = 4, RS amplitude = 2, GT latency = 0, GT amplitude = 0, OSS latency = 2, OSS amplitude = 4. Hence, the GT predictor was selected as the most appropriate predictor and used for all further analyses. All the individual TRFs for the click ERPs and the TRFs are provided in supplementary figures S1-S6. Visual inspection of these individual TRFs confirm that the GT predictor resulted in less noisy TRFs with clear level-dependent wave V peaks. Overall, our results show that level-dependent changes in wave V peaks can be detected in almost all participants, especially when comparing the largest changes in stimulus intensity (72 vs 36 dBA).

### Subcortical responses are consistent across click and speech stimuli

The estimated click ERPs and GT TRFs were compared on a single participant level to investigate the consistency of subcortical responses to clicks and speech. The click ERP and GT TRF wave V peak amplitudes and latencies were significantly correlated across participants and intensity levels (*p <* 0.001, see Fig. 3 leftmost column). To further investigate the level-dependent change in wave V, linear regression lines for the latencies/amplitudes vs. intensity level data were fitted for each participant. The fitted slopes and intercepts for click ERPs and speech TRFs were then correlated across participants (see two right columns of Fig. 3). The latency slopes and intercepts from the GT TRFs were significantly correlated with the click ERP slopes (Pearson’s *r* = 0.57*, p* = 0.02) and intercepts (*r* = 0.83*, p <* 0.001), but this was not the case for amplitude slopes (*r* = 0.29*, p* = 0.3) and intercepts (*r* = 0.16*, p* = 0.5). Overall, these results indicate that level-dependent changes in wave V latencies are consistent across click ERPs and speech TRFs at an individual participant level, but it is unclear if the same applies for the wave V amplitudes. Further work is required to investigate these latency-intensity and amplitude-intensity functions, perhaps using stimuli with more fine-grained click and speech intensity levels.

**Figure 3:**
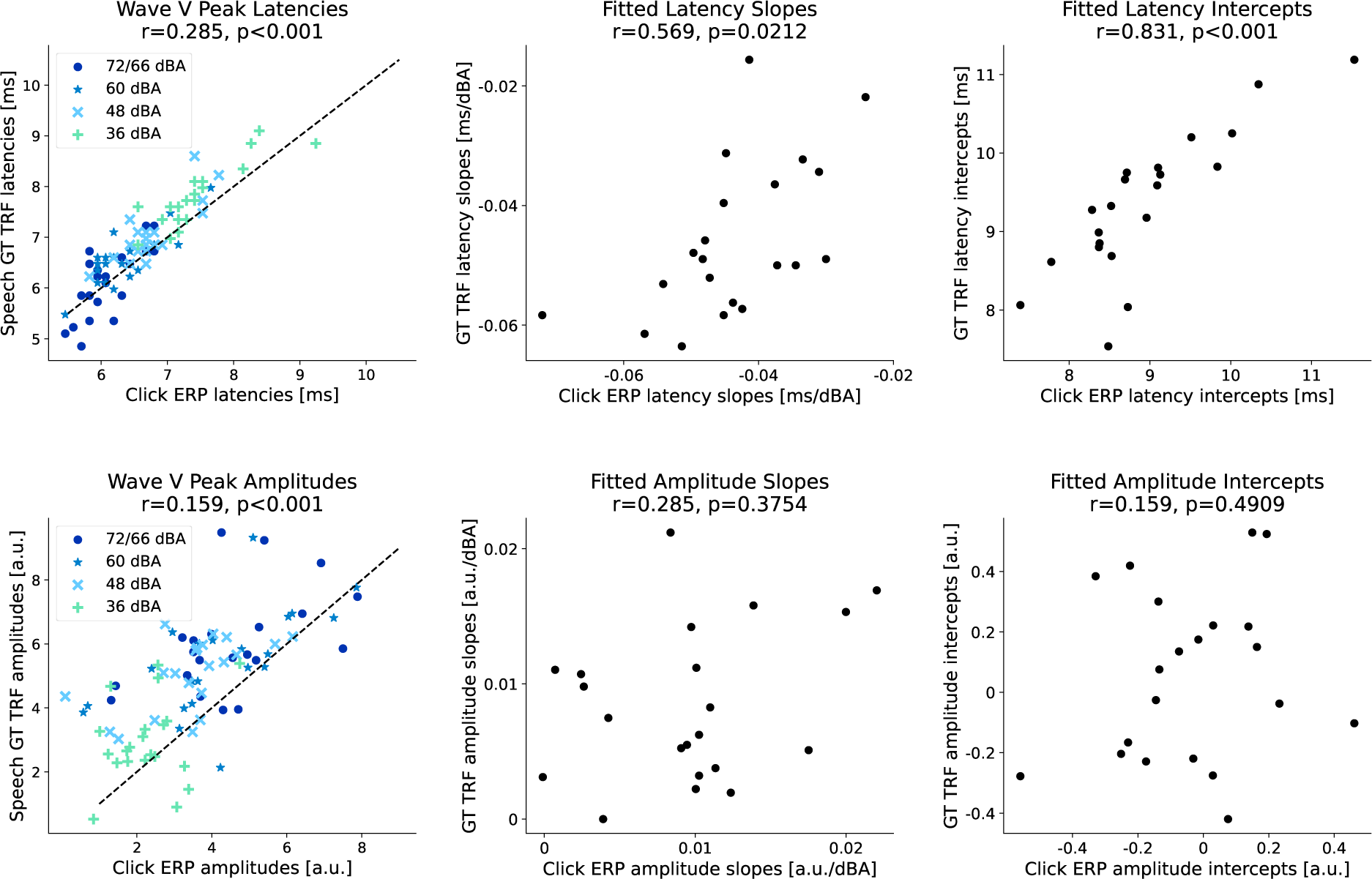
Individual click ERP and speech GT TRF wave V peak amplitudes and latencies. Scatterplots with the individual wave V peak latencies (top row) and amplitudes (bottom row) are shown. Click ERPs are shown on the x-axis and GT TRFs on the y-axis. The leftmost column shows the data for all 4 intensity levels (different marker types are used for each level). The click ERPs and speech TRFs are significantly correlated (Pearson correlations and Holm-Sidak corrected p-values are shown in the subplot titles). Next, linear regression lines were fit using the latency/amplitude vs intensity level data for each participant. The middle column shows the slopes of the regression fits for each participant, and the right column shows the intercepts. The fitted latency slopes and intercepts are significantly correlated across click ERPs and GT TRFs, while the fitted amplitude slopes and intercepts are not (although a weak trend can be observed). These results indicate that the level-dependent changes are somewhat consistent across click ERPs and speech TRFs at a single participant level.

### Level dependent changes in subcortical TRFs to speech stimuli with both long and short durations of fixed intensity

Next, the data from the two experimental conditions were separated. Each condition consisted of 40 minutes of speech stimuli with long (1 minute) and short (5 seconds) durations where intensity was varied across the four possible levels. GT TRFs were estimated in the same manner as before, but separately for the two conditions, and are shown in Fig. 4. TRFs from individual participants for both conditions are shown in supplementary figures S7, S8. It is important to note that each condition has half the amount of data (40 minutes) compared to the results reported in the previous sections (80 minutes), and thus the estimated TRFs are noisier. There were more participants who did not follow the expected trend when comparing peak amplitudes and latencies at 72 vs 36 dBA (long condition amplitudes: 2 participants, latencies: 0 participants; short condition amplitudes: 1 participant, latencies: 1 participant). Neverthe-less, a large majority of participants showed the expected level-dependent changes in wave V amplitude and latency. These results indicate that it is possible to detect level dependent TRFs even when speech intensity changes within relatively short periods that may be comparable to some of the most extreme intensity dynamics in natural speech (e.g., very loud speech changing to whispered speech within a few sentences).

**Figure 4:**
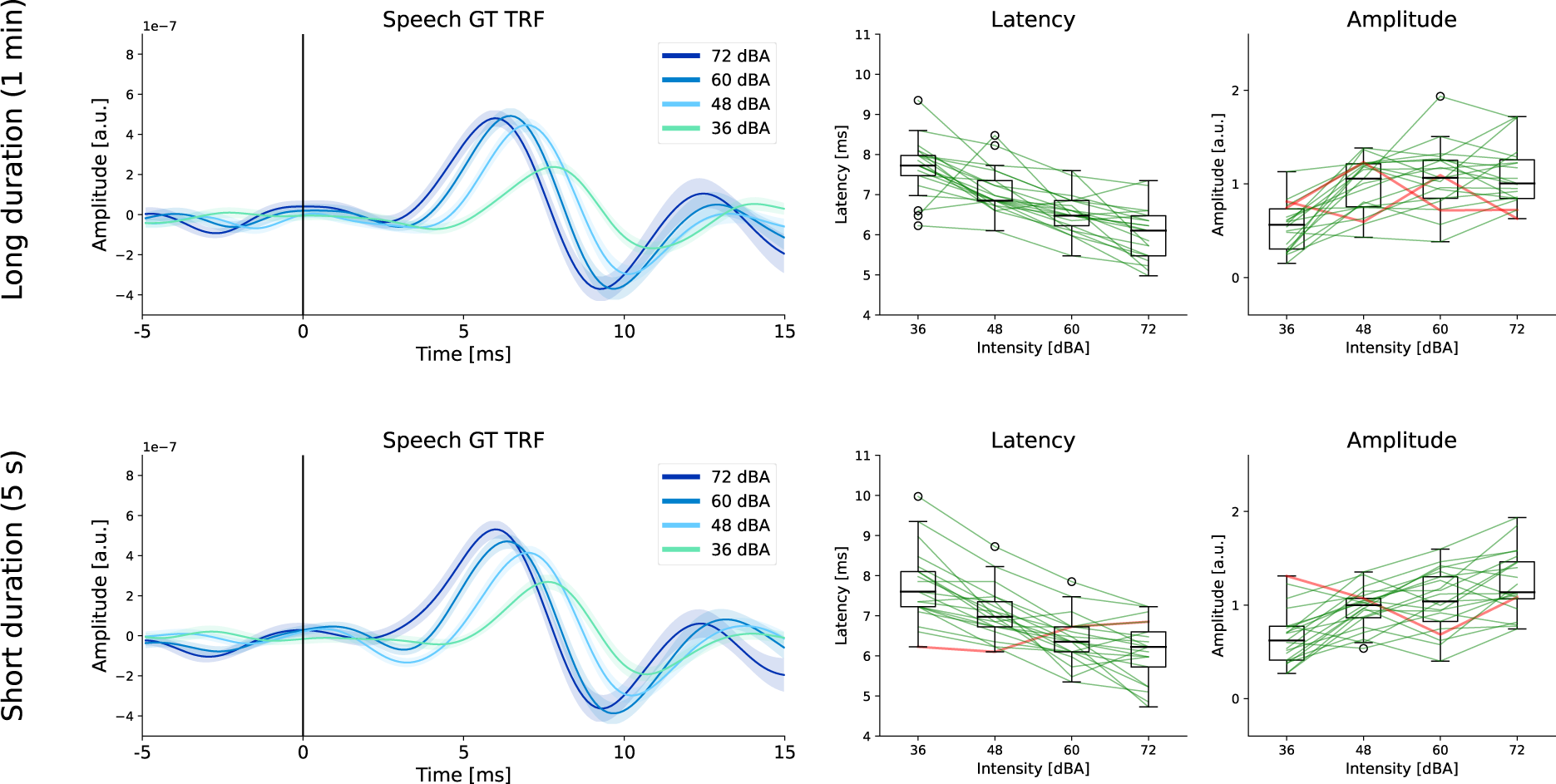
Level dependent TRFs for speech stimuli with long and short durations of fixed intensity. **Top row**: Average TRFs, wave V peak amplitudes and latencies are shown for stimuli with long durations of fixed intensity (1 min changes). **Bottom row**: TRFs and wave V amplitudes and latencies for stimuli with short durations of fixed intensity (5 seconds). Note that there are only 40 minutes of data in each case (10 minutes per level), and this results in noisier TRFs. Even so, level-dependent changes are still clearly seen in a large majority of participants for both long and short duration conditions.

### Amount of data needed to estimate reliable level-dependent TRFs

It is important to determine the amount of data needed to estimate reliable level-dependent TRFs. We used subsets of the data from the short duration fixed intensity condition to estimate TRFs for increasing amounts of data (4, 8, 16 …, 40 minutes). This resulted in 1 to 10 minutes of data within each level for estimating TRFs. To investigate whether the fitted TRFs displayed level-dependent patterns, the wave V peak amplitudes and latencies were extracted for each level, and individual slopes were estimated using a linear fit across the 4 intensity levels, similar to those shown in Fig. 3. The resulting slopes vs. the data length are shown in Fig. 5. Critically, almost all participants show clear level dependent patterns (i.e. negative latency slopes and positive amplitude slopes) after only 5 minutes of data within each level (or 20 minutes of total data). Additionally, we investigated the amount of data required to fit TRFs with reliable wave V peaks within each level using the wave V SNR as a performance metric, similar to prior work (Shan et al., 2024; Kulasingham et al., 2024; Bachmann et al., 2023; Maddox & Lee, 2018; Polonenko & Maddox, 2021). Given that wave V peak amplitudes are larger for high compared to low intensities, it is reasonable to expect that wave V SNR will also be larger at higher intensities (see Fig. 5). As expected, higher intensities have higher wave V SNRs. Critically, our results indicate that almost all participants had a wave V SNR above 0 dB for the highest intensity, with only 6 minutes of data within that level (24 minutes of data in total). This could inform future experimental designs and algorithms that prioritize louder segments of speech to efficiently detect subcortical responses.

**Figure 5:**
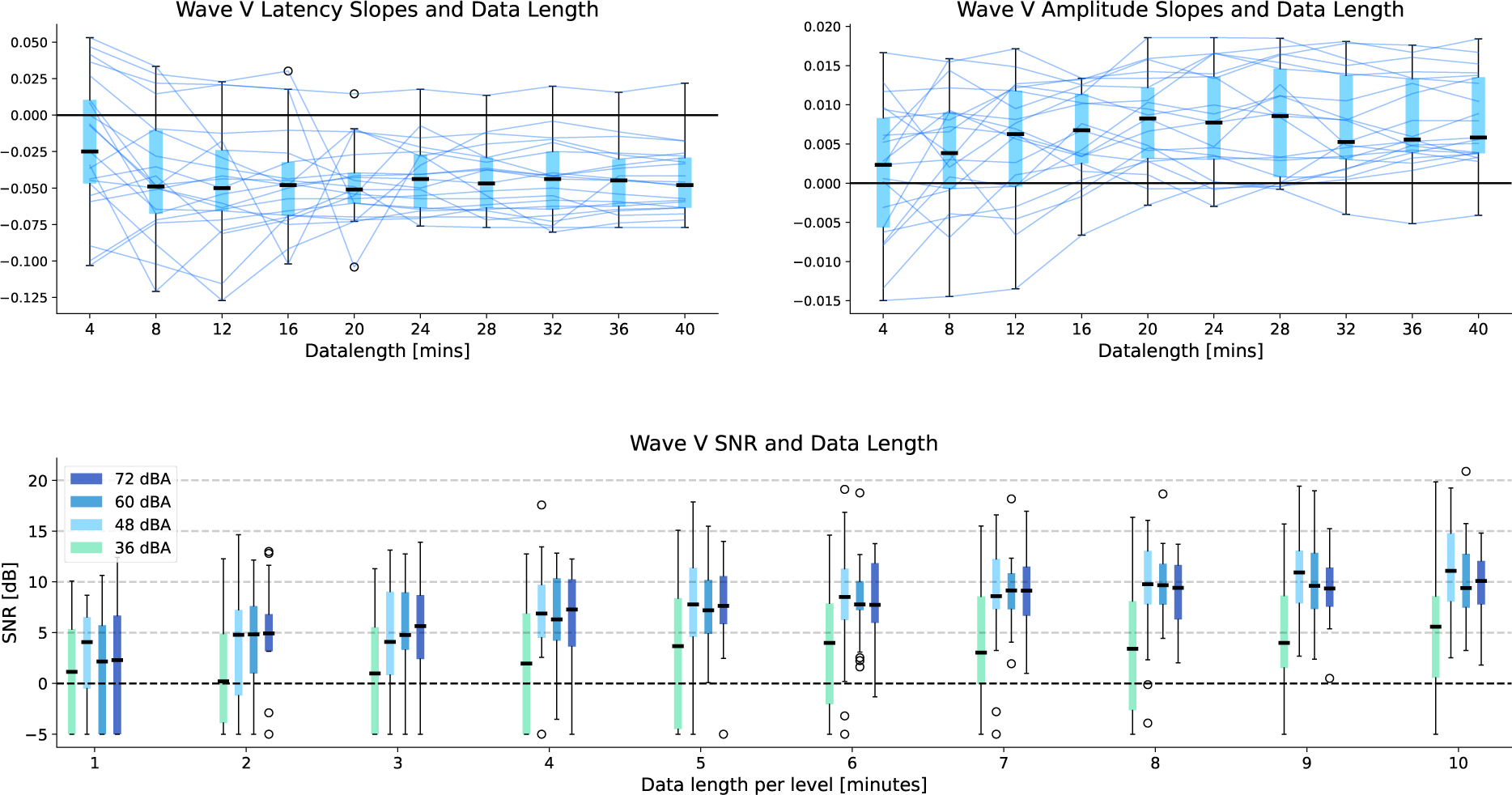
Impact of data length on level-dependent TRFs. Top: Boxplots of the fitted slopes for wave V peak amplitude/latency vs. intensity levels are shown across participants (for the GT TRF) for each data length. A negative slope for latency and a positive slope for amplitude would indicate a level-dependent pattern. Each line represents an individual participant. Almost all participants show level-dependent effects in both latency and slopes after around 5 minutes per level (20 minutes of total data in the jointly fit TRF model). **Bottom:** Boxplots of the wave V SNR across participants (for the GT TRF) are shown for each intensity level and each data length. Around 6 minutes of data within a level (24 minutes of total data) is enough for almost all participants to have wave V SNRs above 0 dB for the loudest intensity. However, some participants are still below 0 dB for the lowest intensity, even with 10 minutes of data within each level.

### Level-dependent TRFs for inherent intensity changes in natural continuous speech

Finally, we investigated whether level-dependent TRFs could be estimated directly from the inherent intensity fluctuations in natural speech, without knowledge of the externally imposed intensity levels. The GT predictor was smoothed using a Hamming window of width 300 ms and used as a coarse measure of intensity level. This intensity measure was then used to split the data into 8 intensity levels, each having an equal amount of data (see Methods for further details). This method would categorize as low intensity both the speech from the trials presented at a low intensity level and the inherently low intensity parts of a trial presented at a high intensity. This could therefore result in a more accurate and fine-grained measure of the actual intensity of the presented stimuli.

The GT TRFs estimated from all 80 minutes of data (both short and long duration conditions) using the above-mentioned method are shown in Fig. 6. Clear level dependent effects are seen in both the average TRF and the individual peak latencies and amplitudes. The latency of the lowest intensity has a larger variation across participants, which could be due to the fact that the peak was too small and noisy for reliable automatic peak detection. The individual TRFs are shown in supplementary figure S9 and confirm that the lowest level does not have a clear peak for many participants. It is noteworthy that level dependent changes are seen for more than 4 levels, indicating that these changes are not only due to the artificially imposed 4 levels of intensity manipulation, but also due to inherent intensity fluctuations in the speech.

**Figure 6:**
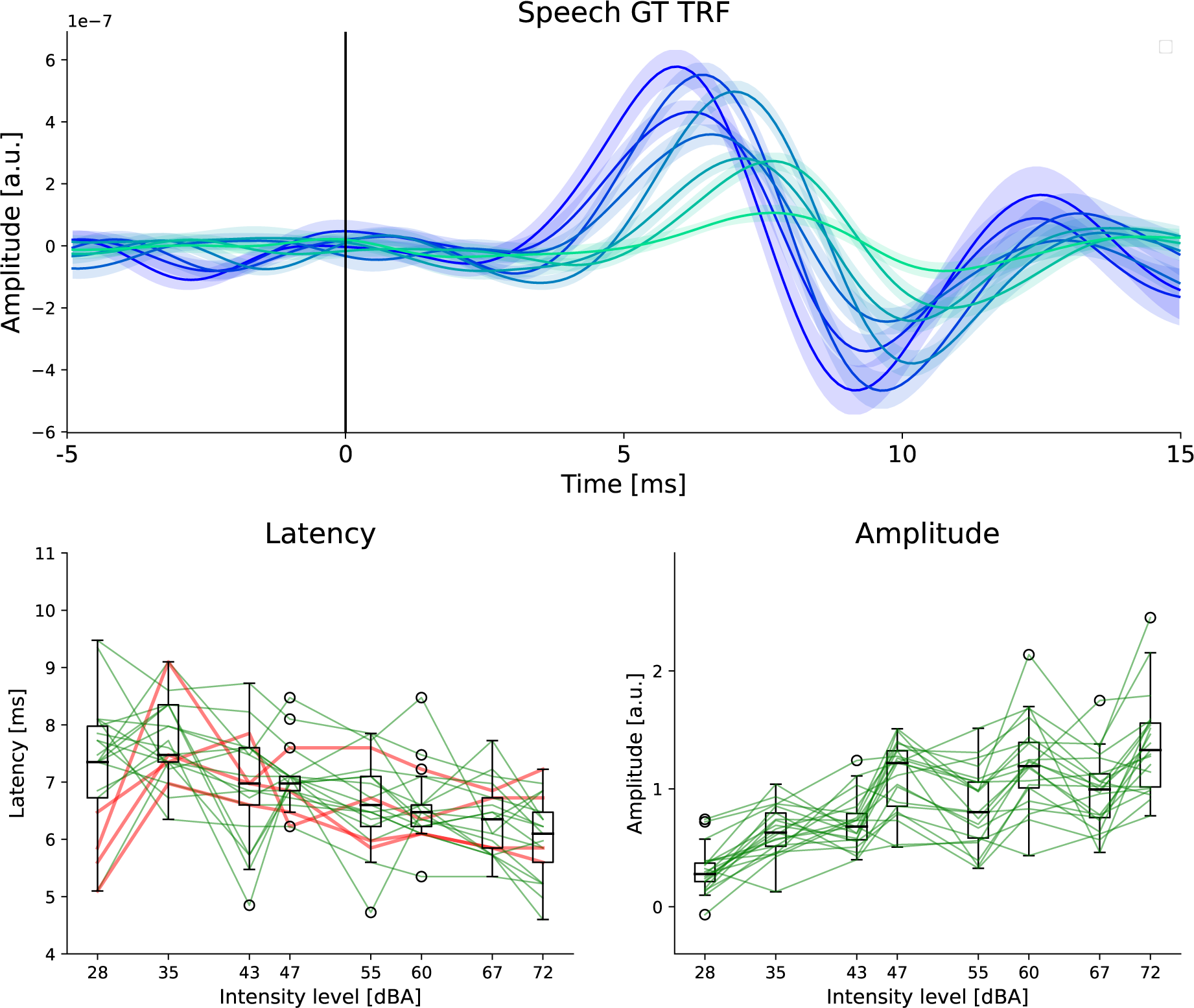
Level dependent TRFs for inherent intensity changes in speech. **Top**: Average GT TRFs and s.e.m. across participants for 8 levels are shown. The darker colors are for higher intensities. **Bottom**: Wave V latency and amplitude changes are shown for 8 levels. Note that the intensity bins were selected to have an equal amount of data in each level, and hence are not equidistant on the log scale. The change between the highest and lowest levels is shown for each participant using red and green lines, following a similar approach to previous figures. Clear level-dependent changes that follow the expected trend (decreasing latency and increasing amplitude, with increasing intensity) are seen for a majority of participants. The latencies at the lowest levels are more variable across participants, which could be due to the fact that the TRFs at the lowest levels are much noisier, and some participants do not have clear wave V peaks.

These results serve as an initial proof-of-concept that the smoothed GT predictor can be used to estimate level dependent TRFs based on the inherent intensity fluctuations in natural speech, and possible alternative measures of inherent intensity level are also explored in the Discussion.

## Discussion

### Level dependent changes in subcortical wave V for continuous speech stimuli

In this work, we developed an experimental paradigm and analysis methods to detect level-dependent changes in subcortical TRFs to continuous speech. The main results of our study correspond to the research questions that were explored and are as follows. 1) Level dependent patterns were detected in subcortical TRFs, with wave V peak latencies decreasing and amplitudes increasing with increasing stimulus intensity. 2) The gammatone filterbank predictor, which does not account for level-dependency in the periphery, was found to be the most suitable to estimate level-dependent TRFs. 3) Level-dependent changes were seen in almost all individuals 4) These level-dependent changes were consistent with level dependent changes in click-evoked wave V peaks at an individual level, especially with regards to peak latency. 5) Level-dependent patterns could be detected in almost all participants with only 20 minutes of total data. Higher intensities of speech required less data to estimate clear wave V peaks (around 6 minutes within each level). 6) Level-dependent TRFs could be detected for stimuli with both short (5 s) and long (1 min) durations of fixed intensity. 7) Finally, we provide a proof-of-concept that level-dependent changes in subcortical TRFs can be detected for the inherent spontaneous fluctuations in stimulus intensity, paving the way for detecting level-dependent subcortical TRFs in natural speech without artificially manipulating stimulus intensity.

### Suitability of auditory models for generating predictors for level-dependent TRFs

Previous work has investigated the suitability of peripheral auditory models for generating predictors for subcortical TRFs, but not with regards to detecting level dependent effects (Kulasingham et al., 2024; Shan et al., 2024). In this work, we compared the RS predictor with predictors derived from peripheral models with adaptation (OSSA, ZIL) and without adaptation (GT, OSS). Although the models that include adaptation were previously found to perform better at estimating wave V peaks for fixed stimulus intensities (Kulasingham et al., 2024), they did not result in clear level dependent latency effects in the estimated TRFs.

This surprising result can be explained by the fact that many of these models account for nonlinear effects of level-dependency when generating their outputs. When these model outputs are subsequently used as an input predictor, the effect of stimulus intensity is thereby ‘linearized’ in the TRF estimation problem. This results in two outcomes: a better linear TRF model fit, and a lack of level-dependency in the estimated TRFs. Although the former is desirable for some applications (e.g., detecting wave V peaks with less data), the latter is detrimental for our goal in this work, which is to detect the level-dependent changes in TRF wave V, analogous to level-dependent changes in the click ERPs. Therefore, these more complex models with adaptation (OSSA, ZIL) were not suitable, and simpler filterbank based models (GT, OSS) were preferred. Of the latter two, we found that the GT predictor performed better, with more participants showing a clear level-dependent effect. Even though the use of the GT predictor likely resulted in noisier TRFs compared to the ZIL and OSSA models, we still observed clear level-dependent effects in wave V peaks in nearly all participants. Therefore, we conclude that, of the 5 predictors that were investigated, the GT predictor is the most suitable for estimating level dependent subcortical TRFs. Future work could investigate modifications to these models or other models (e.g., (Verhulst et al., 2015)) that may be able to leverage the desirable properties of complex models (such as higher SNR wave V peaks), while still retaining level dependent effects in the TRF.

### Methodological considerations when estimating level-dependent TRFs

Level-dependent TRFs were estimated using a regression model with binned predictors, similar to prior work with cortical TRFs (Drennan & Lalor, 2019; Lindboom et al., 2023). The predictors for each level should be jointly estimated in a combined regression model, since separate estimation may lead to spurious effects caused by collinearities across the predictors, especially for the inherent intensity analysis, where the intensity bins could change rapidly. An additional consideration is that the amount of data in each predictor (i.e., the amount of data in each intensity level bin) can impact the resulting TRFs. Intensity bins with lower amounts of data can result in noisier TRFs, which may also result in incorrect wave V peak amplitudes or latencies. In our experiment, we constructed the stimuli with an equal amount of data in each intensity bin. On the other hand, for our preliminary investigation on level-dependent TRFs from inherent intensity fluctuations, we defined the intensity bins to have an equal amount of data. This resulted in intensity bins that were not equally spaced, but did allow for balanced estimation of the TRF in each bin.

In this work, the methods were chosen (e.g., smoothing window size) to reliably detect a wave V peak in the most participants, since this was the primary metric that was used to evaluate level-dependency. However, earlier waves of the ABR also provide information about the function of different stages of the auditory pathway. We could not see clear early waves in subcortical TRFs, which is not surprising given prior work (Maddox & Lee, 2018; Kulasingham et al., 2024; Bachmann et al., 2023) and the estimation methods used in our study. Future work is needed to determine if such waves can be reliably detected in subcortical TRFs.

Finally, in this work we did not use any form of regularization, although this is commonly done for cortical TRF studies. We chose to not use regularization in order to follow the same methods as prior work on subcortical TRFs (Maddox & Lee, 2018; Polonenko & Maddox, 2021; Kulasingham et al., 2024) and to simplify the analysis pipeline by avoiding the optimization of a regularization parameter. Additionally, regularization could lead to confounds in the amplitudes and latencies of level-dependent TRF peaks. Our results indicate that regularization is not necessary for subcortical TRFs, at least for the data lengths used in this work, but may be useful in other situations.

### Estimating level-dependent TRFs for inherent intensity changes in speech

The TRF method could be extended and applied to investigate level-dependent subcortical responses to the inherent intensity fluctuations in natural speech. Our proof-of-concept results show that it is possible to detect these changes by selecting intensity bins based on a smoothed version of the GT predictor. We used a Hamming window with a width of 300 ms to smooth the GT predictor, to capture fast changes in intensity dynamics near the syllable rate, while still allowing for reasonably slow changes from one intensity bin to another. Alternatively, a different window size or a different measure of intensity, perhaps directly from the speech stimulus itself, could be used instead, and may improve level dependent TRF results. A more systematic investigation of the optimal smoothing window or intensity measure was not explored here since it is beyond the scope of this work.

Another caveat when estimating TRFs based on the inherent intensity fluctuations in natural speech is that the lower intensity bins may have different acoustic properties compared to higher bins (e.g., fricatives at lower intensities vs vowels at higher intensities). Therefore, differences in the corresponding TRFs estimated at each bin could be also driven by these other acoustic properties rather than purely by the intensity (Skoe & Kraus, 2010). For example, our analysis of inherent intensity fluctuations had 8 levels, each pair of which roughly corresponded to the 4 stimulus intensities that were used. There seemed to be larger variation of wave V amplitudes within each pair of the 8 levels (see Fig. 6), which suggests that the level-dependent amplitude effect is more pronounced for inherent intensity fluctuations within each of the 4 stimulus intensity levels, possibly due to different speech sounds in each of pair of levels. However, all our other analysesinvolved estimating TRFs based on speech presented at different overall intensity levels, ensuring an even distribution of speech sounds across all intensity bins. These issues must be taken into account in future work when investigating level dependent TRFs to inherent intensity fluctuations in natural speech.

### Possible applications in clinical audiology and hearing technology

Understanding level-dependent changes in TRF wave V peaks is essential to be able to compare sub-cortical TRFs to conventional ABR ERPs, which have been extensively studied in scientific and clinical contexts. Investigating level-dependent subcortical responses to continuous speech is important for clinical applications such as determining hearing thresholds in a comfortable and ecologically relevant environment (Hamilton & Huth, 2020; Lunner et al., 2020). Additionally, hearing aids often filter out unnatural transient stimuli such as clicks, and are hard to use in conjunction with conventional ABR ERP protocols. Therefore, subcortical TRFs to continuous speech may provide an objective measure to test and calibrate hearing aids and investigate the impact of hearing aid speech enhancement algorithms on the early auditory system. Finally, developing an algorithmic pipeline for detecting level-dependent subcortical responses to the inherent intensity fluctuations in speech could lead to future adaptive ‘neuro-steered’ hearing aids (Geirnaert et al., 2021) that adjust speech enhancement algorithms and gain based on dynamically detected subcortical responses to natural speech.

## Supplementary Figures

**Figure S1:**
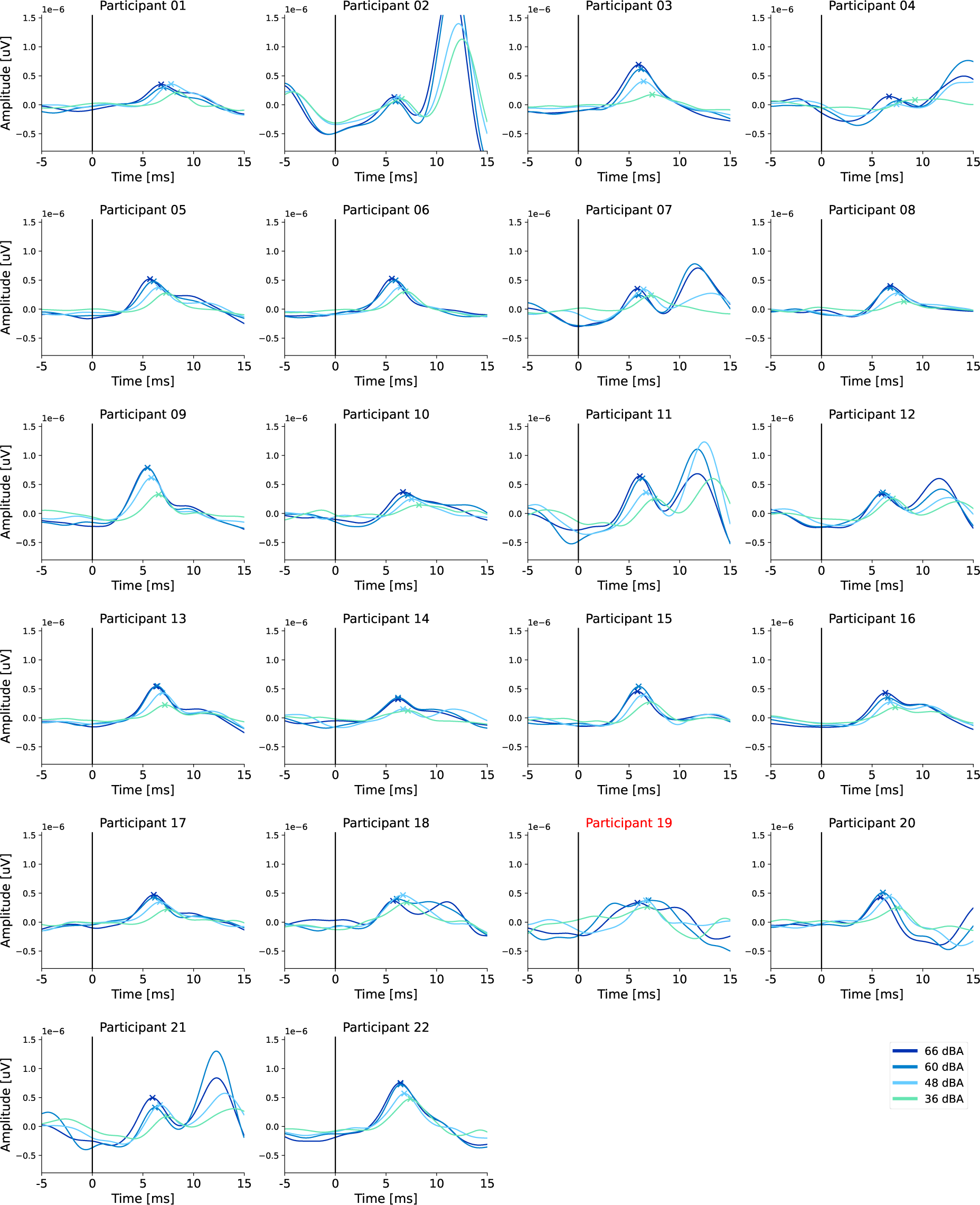
Individual click ERPs. All the click ERPs for each participant calculated on the full dataset are shown. Some participants also have a large post-auricular muscle reflex or middle latency responses. The peak in the range of 4-10 ms is automatically detected as the wave V peak (marked with an X in the plots). Participant 19 was rejected and not analyzed due to the lack of clear click ERPs or speech TRFs, but is shown here for completeness.

**Figure S2:**
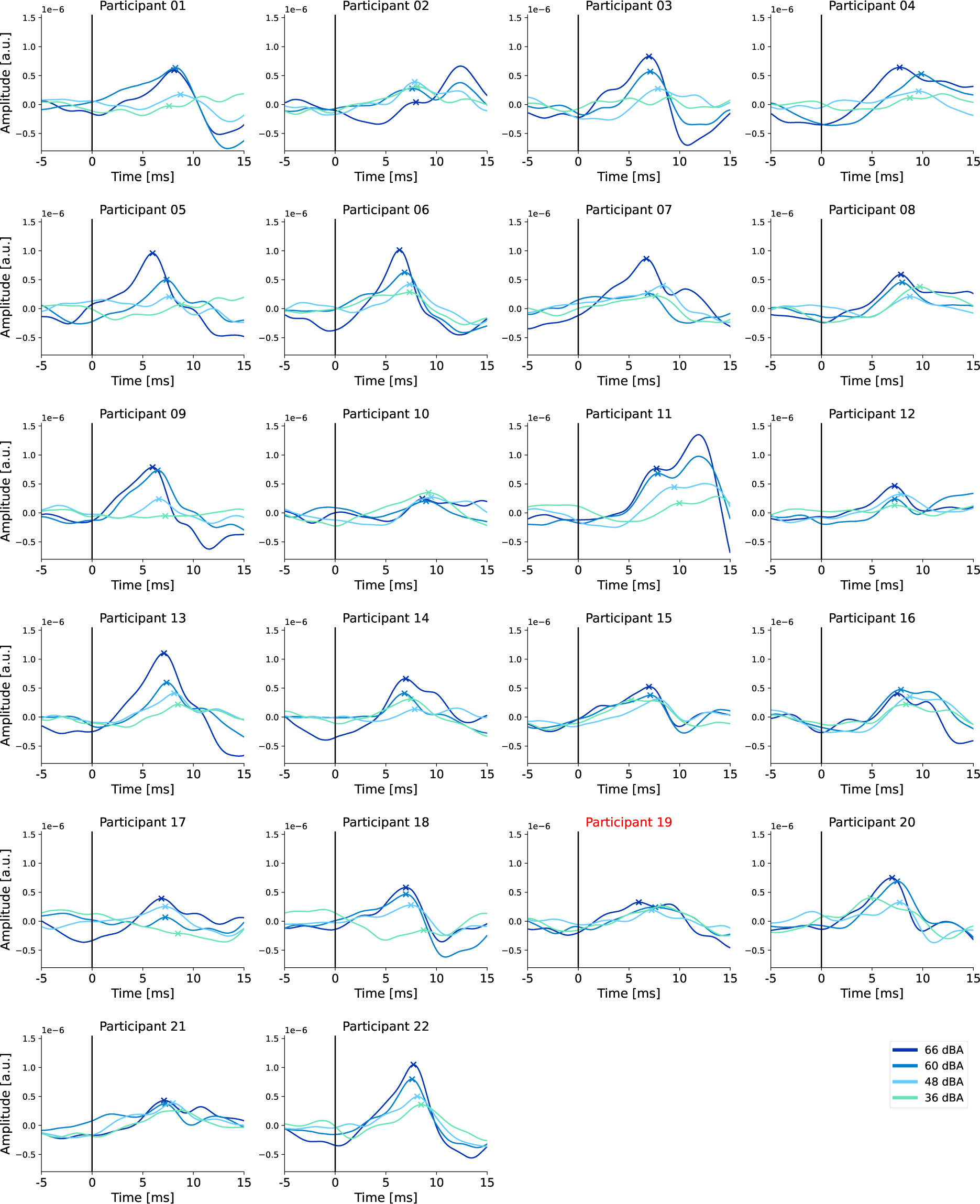
Individual Speech TRFs: RS Predictor. RS TRFs for each participant calculated on the full dataset are shown. Note that TRFs are quite noisy and have broad peaks.

**Figure S3:**
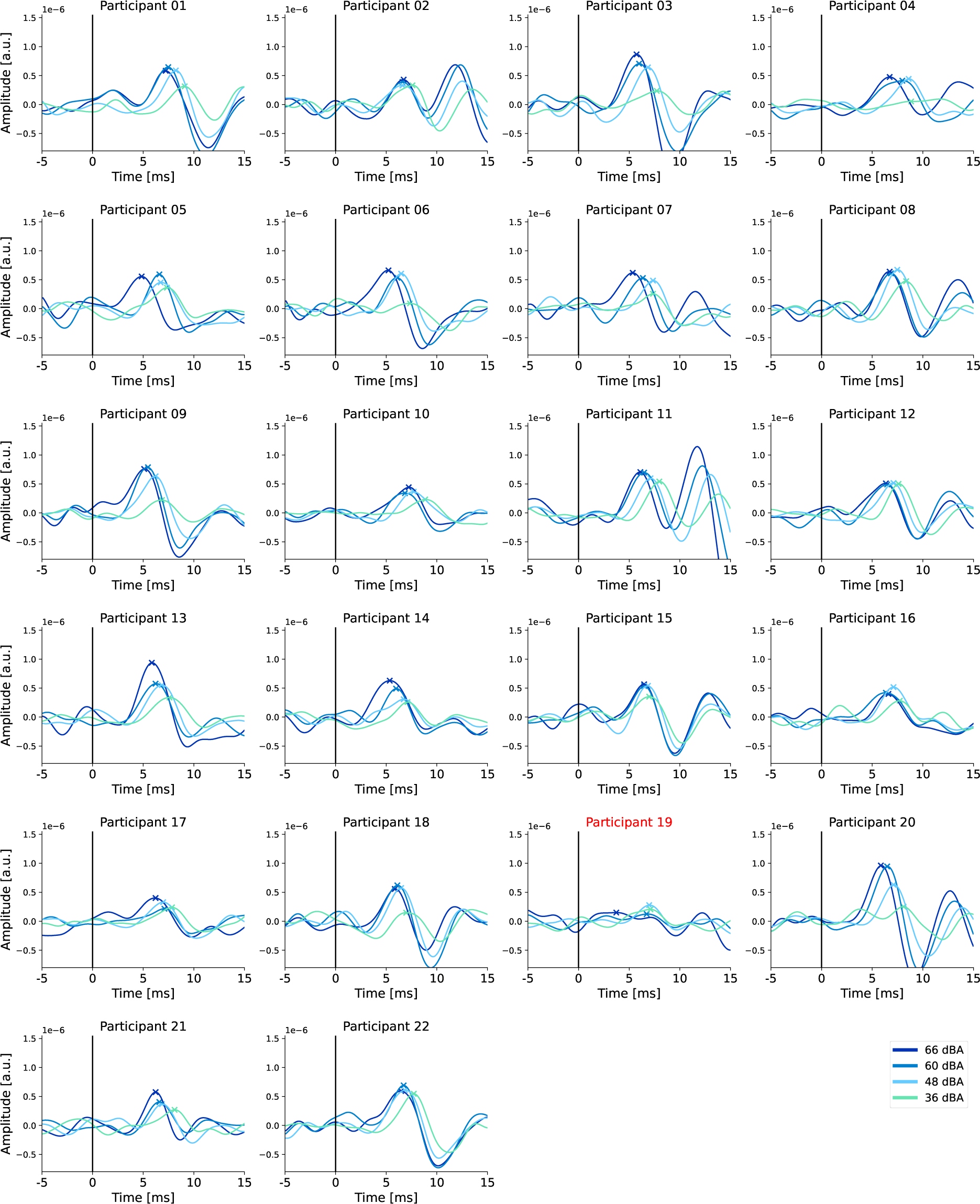
Individual Speech TRFs: GT Predictor. GT TRFs for each participant calculated on the full dataset are shown. Note that both level-dependent amplitude and latency effects can be seen for most participants.

**Figure S4:**
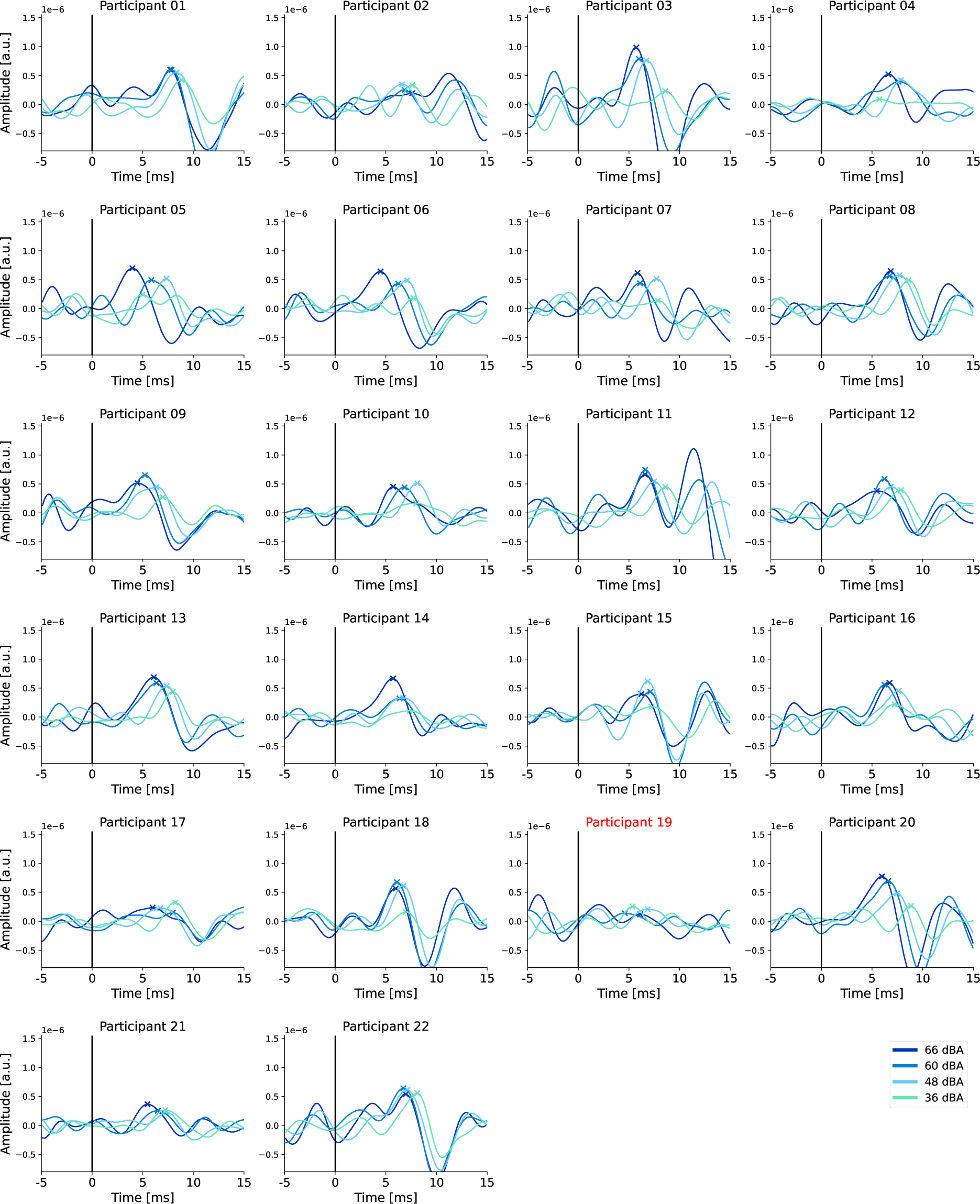
Individual Speech TRFs: OSS Predictor. OSS TRFs for each participant calculated on the full dataset are shown. Note that TRFs are noisier than GT TRFs.

**Figure S5:**
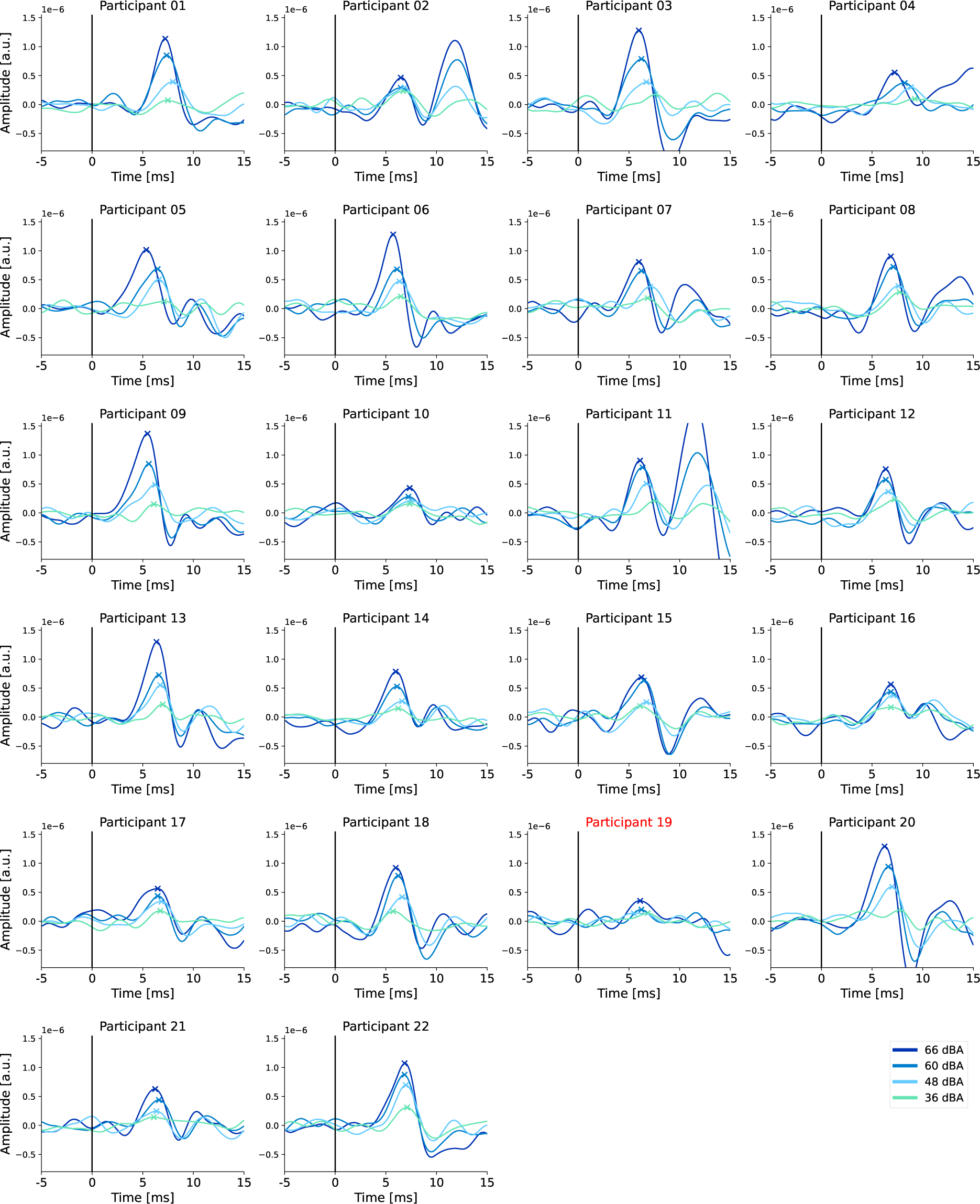
Individual Speech TRFs: OSSA Predictor. OSSA TRFs for each participant calculated on the full dataset are shown. Note that level-dependent latency effects are not as prominent.

**Figure S6:**
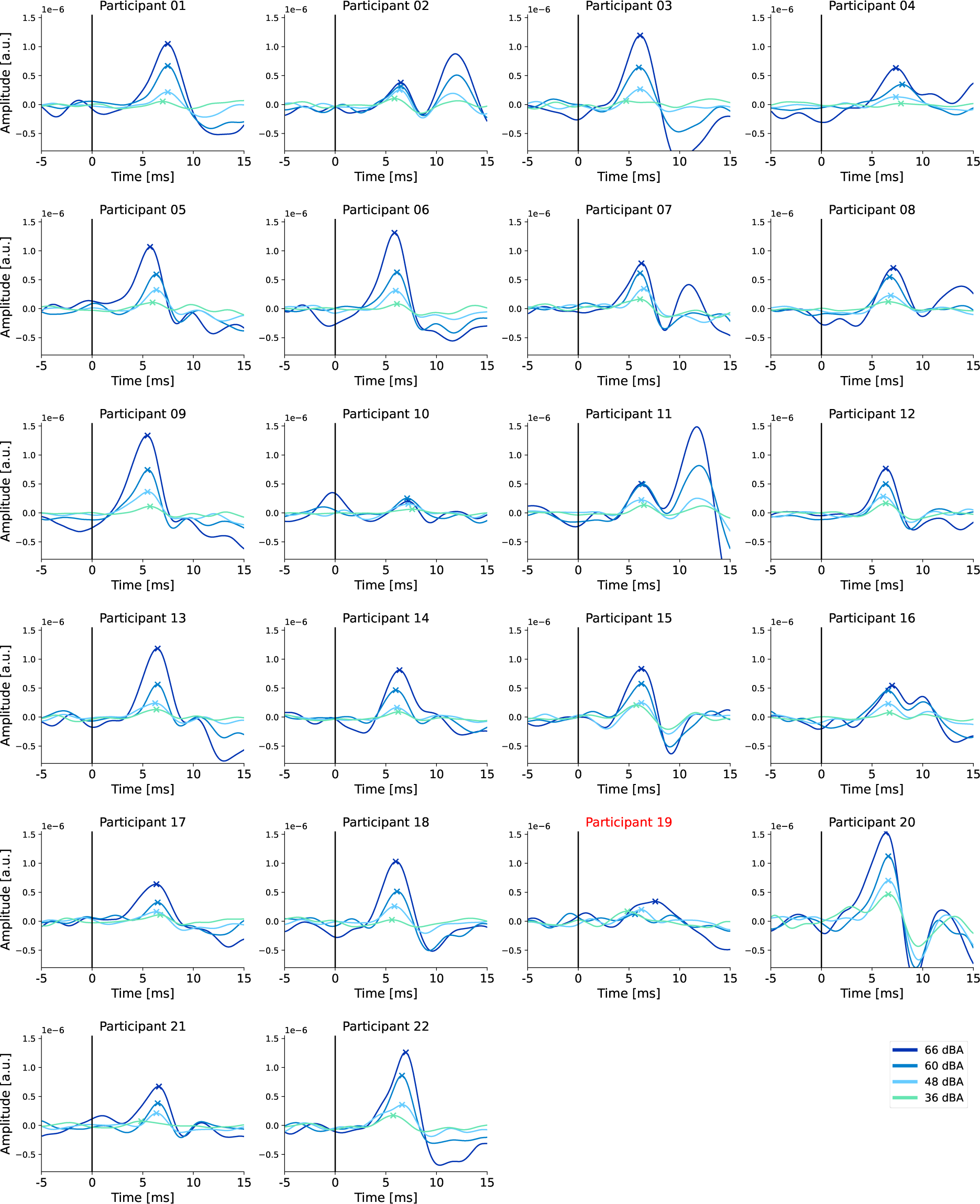
Individual Speech TRFs: ZIL Predictor. ZIL TRFs for each participant calculated on the full dataset are shown. Note that level-dependent latency effects are absent or against the expected trend in most participants.

**Figure S7:**
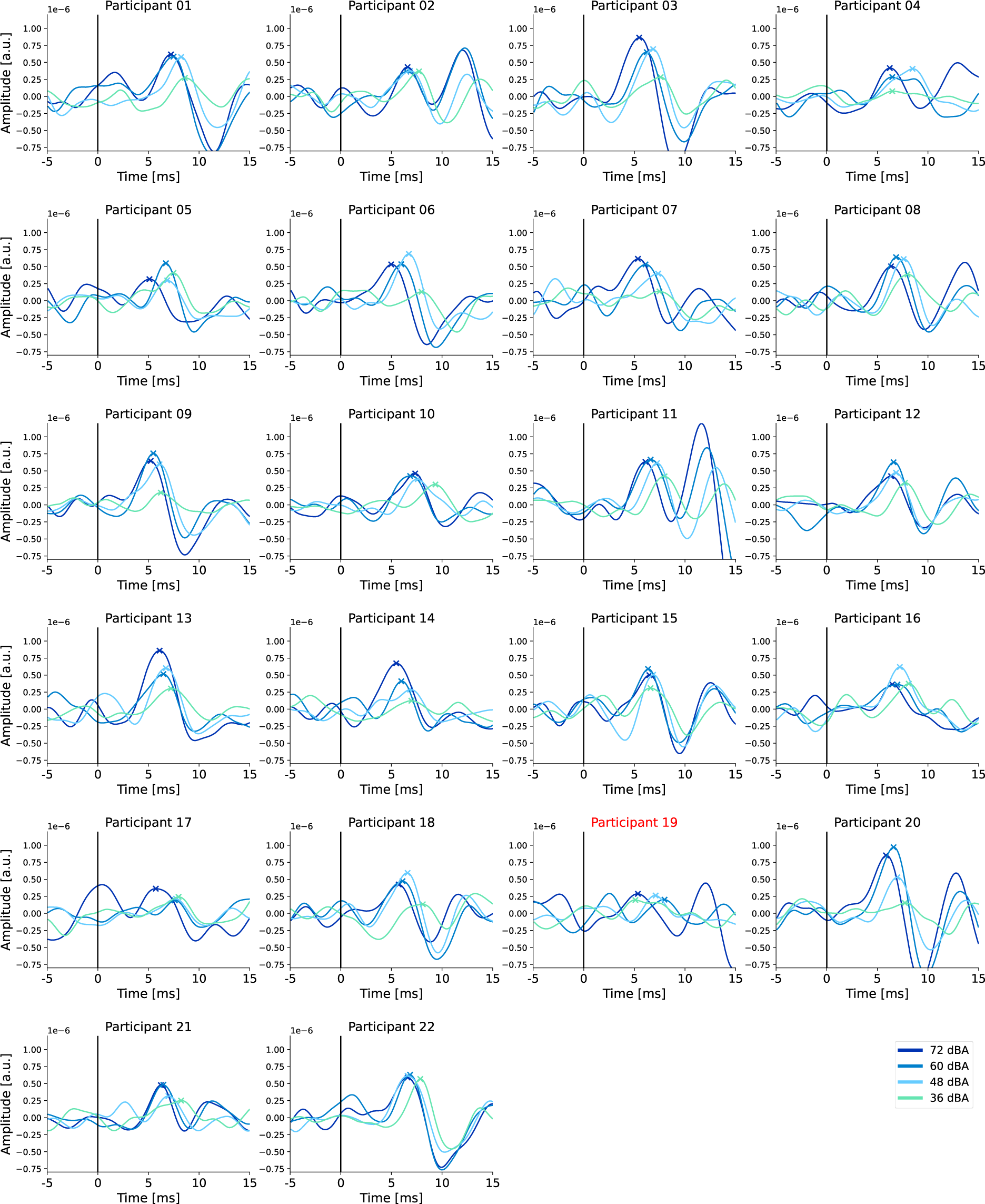
Individual Speech GT TRFs for the long duration fixed intensity condition. GT TRFs for each participant calculated on the 40 minutes of long duration fixed intensity condition are shown.

**Figure S8:**
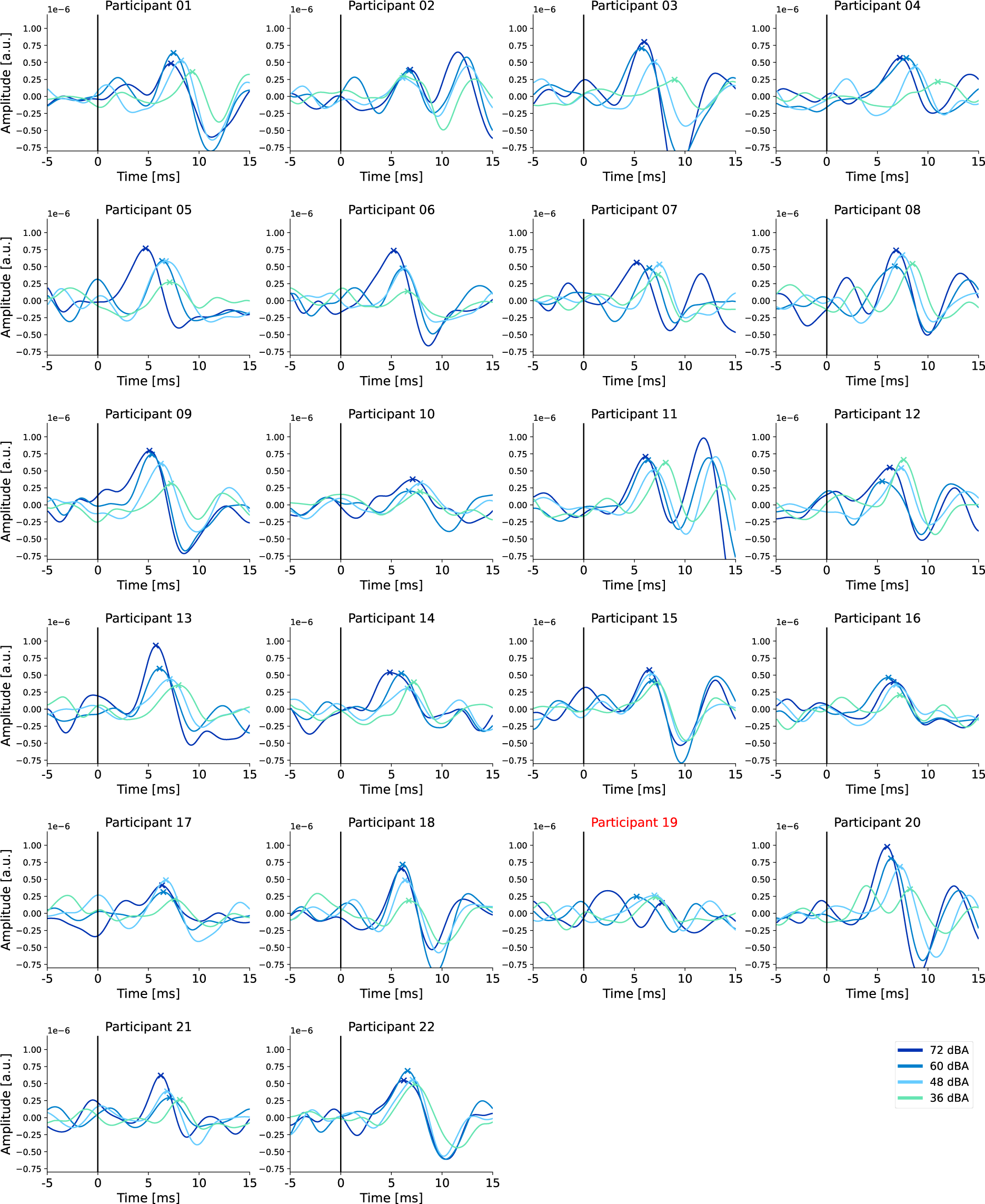
Individual Speech GT TRFs for the short duration fixed intensity condition. GT TRFs for each participant calculated on the 40 minutes of short duration fixed intensity condition are shown.

**Figure S9:**
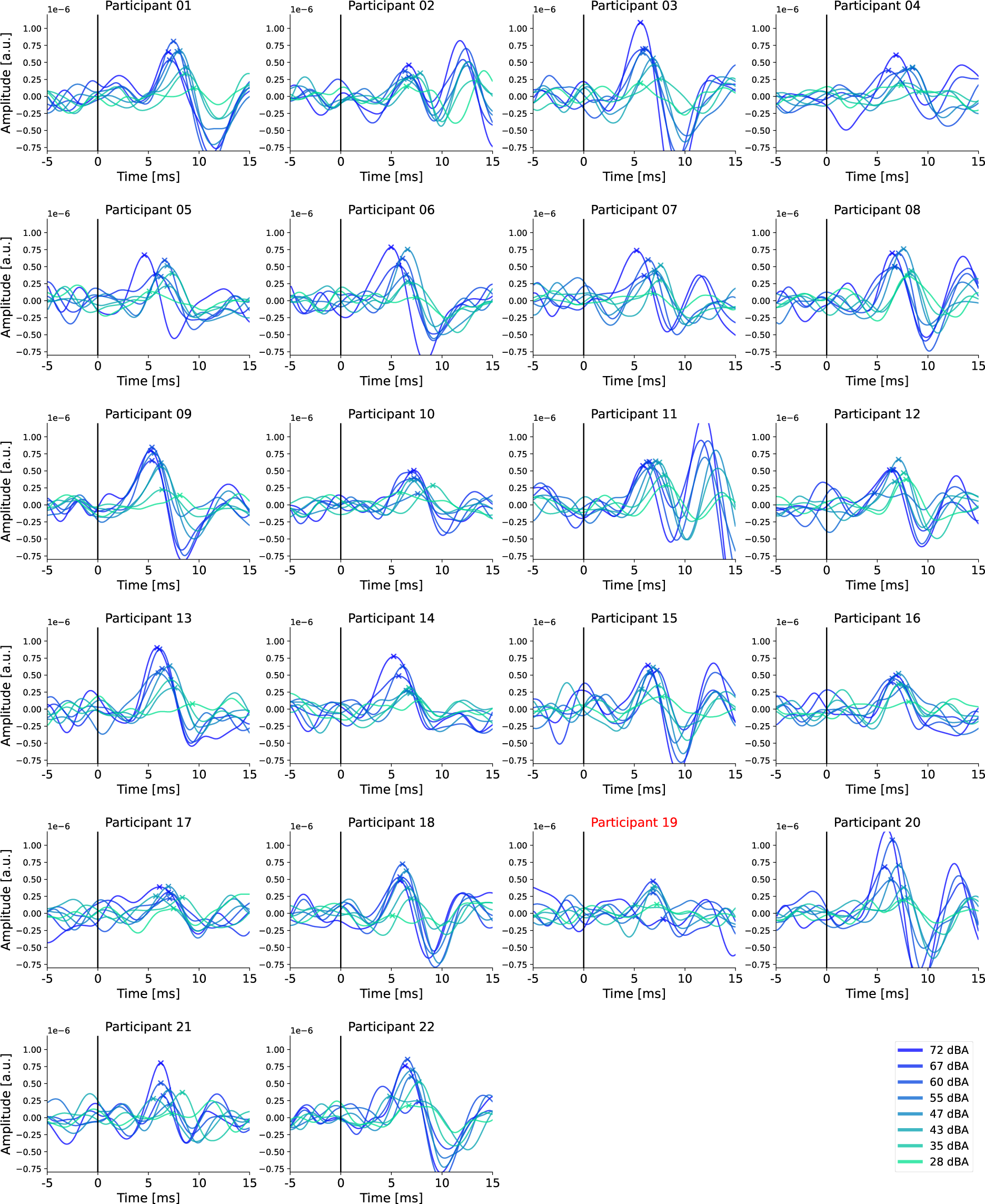
Individual Speech GT TRFs based on inherent changes in intensity level. GT TRFs for each participant calculated for the inherent level changes are shown. Note that most participants have a clear trend of increasing latency with decreasing level.

